# Biofilms with a dash of color: A hands-on activity for school students to build a biofilm model, and use it to understand antibiotic tolerance in biofilms

**DOI:** 10.1101/2022.06.27.497701

**Authors:** Snehal Kadam, Kashish Methwani, Karishma S Kaushik

## Abstract

It is increasingly recognized that microbes, such as bacteria, exist in communities or biofilms, both in the environment and human body. However, school biology curriculum continues to focus on the free-floating form of bacterial life, with minimal descriptions of biofilms. Consequently, there is a need to introduce biofilms to school students, to not only to develop a fundamental understanding of microbial life, but to also highlight the challenges posed by biofilm infections to antibiotic treatment. We have developed a hands-on activity in which students build a biofilm model, and use it to understand the role of the extracellular matrix in the antibiotic tolerance of biofilms. The activity uses simple, easy to obtain supplies, and can be conducted in an in-person or virtual format, with students participating from school or home. We conducted the activity in virtual mode for a group of 59 school students across India, and present feedback and learnings that could be used to execute and adapt this accessible and engaging science experience.

## INTRODUCTION

In contrast to their popular depictions as free-living (planktonic) forms, microbes such as bacteria and fungi most often exist in self-assembled communities or biofilms (1). Responsible for a range of human infections, bacterial biofilms consist of clumps of bacteria embedded in an extracellular matrix (glue), produced by the bacteria themselves (2) (3) While antibiotics successfully eliminate free-living bacteria, biofilms display increased tolerance to antibiotics (4). This is in part due to the presence of the extracellular matrix in biofilms, that limits the penetration of select antibiotics (5). Previous hands-on biofilm activities have developed approaches to introduce important concepts such as the structure and robust nature of biofilms to younger age groups (6) (7) (8). Extending on these previous activities, we present a hands-on activity for school students to build a biofilm model, and use it to understand the role of the extracellular matrix in the antibiotic tolerance of biofilms. Using simple and easy to obtain supplies, this hands-on activity can be conducted in a classroom or via virtual platforms, for students at school or home. We conducted this hands-on activity via virtual mode for 59 students across India during pandemic-related school closures, and present feedback based on the home-based science experience.

## PROCEDURE

### Materials

Each student will need a set of these materials. Following the activity, the model and used materials can be discarded in a regular trash bin.

- Two clear, plastic, disposable glasses
- Transparent or translucent hair gel or face wash or glue (100 mL)
- Sprinkles or small beads or confetti, preferably different colors and shapes (a handful)
- Water (100 mL)
- Gel or liquid food coloring (1 bottle, any color) or any colored liquid (10 mL)
- 2 tablespoons
- Thick marker pen

### Safety issues

There is a small risk of ingestion of hair dye, face wash, confetti, or beads, for which supervision is recommended for young children.

### The hands-on activity

Before the session, the instructor could provide a brief introduction to microbes, biofilms and antibiotic tolerance in biofilms (**Appendix 1 and 2)**.

The step-by-step instructions for the students are **(Figure 1)**:

**Figure 1:**
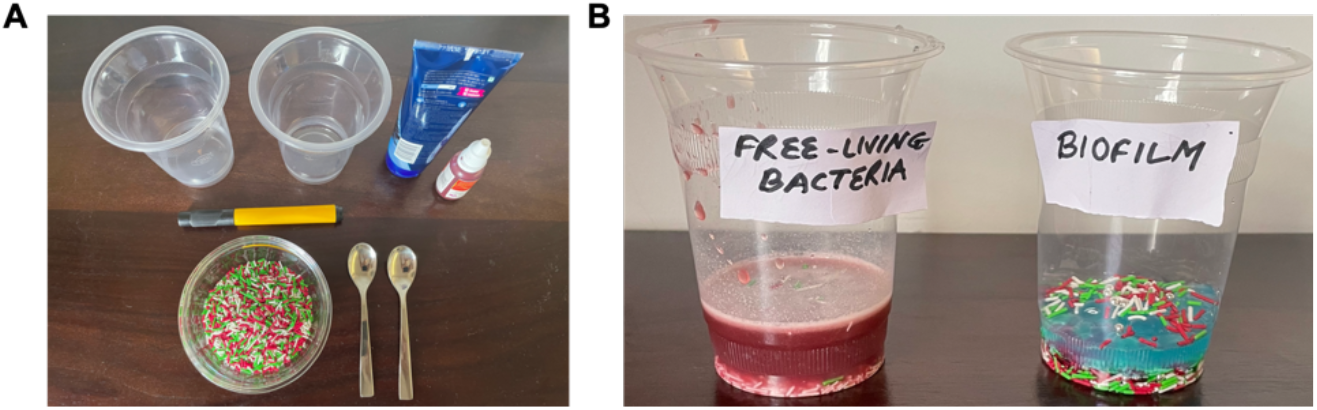
Building a biofilm and free-living bacteria model using simple, easy to obtain supplies. **(A)** To mimic bacteria, students can use sprinkles, glitter, beads, or confetti. To resemble the extracellular matrix, transparent or semi-translucent hair gel or face wash can be used. Food color or any colored liquid can be used to mimic the antibiotic. **(B)** Briefly, students spread ‘bacteria’, across the bottom of a container, and add the ‘extracellular matrix’ in two parts, swirling in between. The free-living bacteria model is built adding water to the ‘bacteria’. The water appears pink due to the color of the sprinkles; this can be prevented with beads or glitter.

- Take the two clear, plastic, disposable glasses, and label one glass ‘biofilm’ and the other ‘free-living bacteria’.
- In both glasses, spread sprinkles or small beads or confetti (represent different types of bacteria), across the bottom of the container.
- In one of the glasses, add half of the hair gel or face wash or glue (represents the biofilm ‘matrix’), swirl to mix, and add more hair gel on top of the swirled mixture. This represents the ‘biofilm’ model.
- In the second glass, add 100 mL of water to the sprinkles, and swirl the mixture. This represents ‘free-living bacteria’.
- Add several drops of food color or gel or liquid (to represent antibiotic) to the ‘biofilm’ and the ‘free-living bacteria’. Do NOT mix the food color, but let it seep on its own.

It is expected that, in the ‘biofilm’ model, the food color (‘antibiotic’) accumulates at the top of the ‘biofilm’, or shows minimal penetration through the ‘extracellular matrix’. In doing so, it has limited access to the ‘bacteria’ embedded in the ‘extracellular matrix’. On the other hand, for the ‘free-living bacteria’, the food color (‘antibiotic’) is expected to penetrate quickly through the water and surround the ‘free-living bacteria’ **(Figure 2)**. In doing so, this mimics the role of extracellular matrix in impeding the penetration of antibiotics through the biofilm, which contributes to the antibiotic tolerance of bacteria in biofilms.

**Figure 2:**
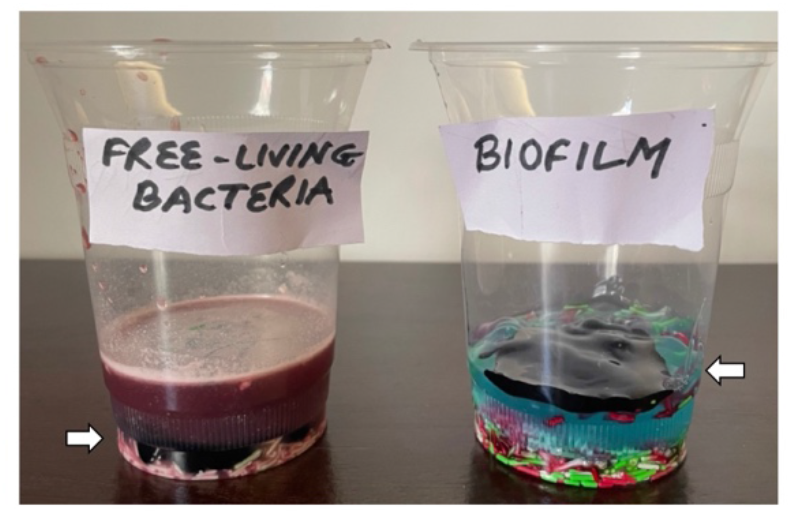
Using the biofilm model to understand antibiotic tolerance in biofilms, in comparison with the model of free-living bacteria. For this, students will add several drops of food color (gel or liquid) to the ‘biofilm’ model and the ‘free-living bacteria’. The food color is used to represent an antibiotic, being used to ‘treat the biofilm’. The expected observation is that, in the ‘biofilm’ model, the food color (‘antibiotic’) accumulates at the top of the ‘biofilm’, or shows minimal penetration through the ‘extracellular matrix’. In doing so, it has limited access to the ‘bacteria’ embedded in the ‘extracellular matrix’. On the other hand, for the ‘free-living bacteria’, the food color (‘antibiotic’) is expected to penetrate quickly through the water and surround the ‘free-living bacteria’.

### Student feedback form the virtual delivery of the hands-on activity

We delivered the hands-on activity in a virtual format to a group of 59 school students from across India with collection of pre-and post-session feedback (**Appendices 3 and 4)**. It is unclear why 7 participants did not provide post-session feedback, or left the session prior to completion. Based on pre-session feedback, only 37% of participants (n=22/59) had heard of biofilms, and even fewer (12%, n=7/59) had done a hands-on activity on biofilms (**Figure 3**). Based on post-activity feedback, majority of the participants were able to grasp the key concepts related to antibiotic tolerance in biofilms, including the extracellular matrix being the major difference between free-living bacteria and biofilms (70%, n=36/52), and the role of the matrix in preventing the penetration of antibiotics (67%, n=35/52) (**Figure 4**). With respect to observations related to the antibiotic response in the model of free-living bacteria, 54% of participants (n=28/52) responded that they observed antibiotic penetration through the model, in contrast to the biofilm model. On the other hand, 46% (n=24/52) responded that they observed antibiotic to not penetrate through the model of free-living bacteria. Based on this, future adaptations of this hands-on activity could focus on clarifying this observation better, maybe by spending some time on the final differences between the antibiotic (food color) penetration in the two models. The majority of participants reported the session to be fun and informative, and responded that they would recommend the hands-on activity. Further, the session itself was hugely interactive (via the chat window) during the content delivery and hands-on component, with the students asking questions about biofilms to the instructor **(Table 1)**, describing their observations with their peers **(Table 2)**, and providing feedback on the experience **(Table 3)**.

**Table 1:** Select feedback from participants with questions related to biofilms asked during the session.

|  |
| --- |
| What are bad biofilms? |
| I could not understand the difference between the good and bad biofilms. |
| On the teeth, does saliva allow bacteria to attach? |
| Can biofilms harm our body or do they simply protect the bacteria? |
| Are biofilms form through which bacteria are becoming antibiotic resistance? |

**Table 2:** Select feedback from participants related to observations of the biofilm model and antibiotic tolerance in biofilms.

|  |
| --- |
| In the first glass the substances are stuck together and in the second one they are living. |
| In the biofilm glass, the color doesn't touch the bottom of the glass. |
| In the free-living bacteria glass, the color mixed and the bacteria are free-living. |
| In the 1st one they are all together in 2nd they are apart. |
| The color doesn't mix with the biofilm. |

**Table 3:** Select feedback from participants related to the experience at the hands-on session.

|  |
| --- |
| It was like really building the home of the microbes. |
| Thank you for the new experiment. |
| It was my best webinar till now! |
| It was a nice session. |
| This class was very interesting. |

**Figure 3:**
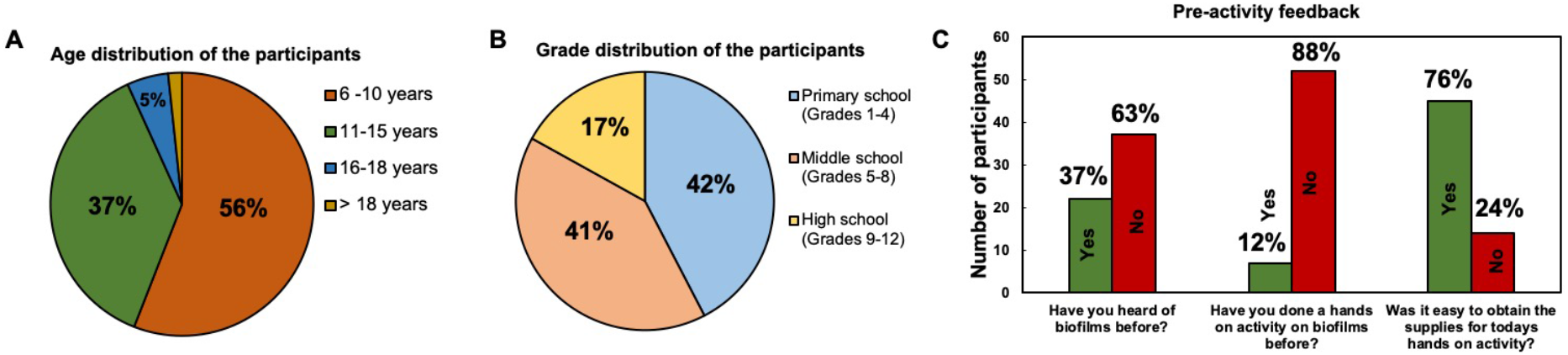
Pre-activity feedback from participants with respect to age, grade, and prior familiarity with biofilms. **(A)** The majority of the participants, 56% (n=33/59) were in the age group of 6-10 years, with 37% (n=/59) ion the age group of 11-15 years. **(B)** With respect to school education level, 42% (n=25/59) of the participants were in primary or elementary school (grades 1-4), and 41% (n=24/59) of the participants were in middle school (grades 5-8). **(C)** With respect to prior familiarity with biofilms, 63% of participants (n=37/59) responded that they had not heard of biofilms before this session, and only 12% (n=7/59) had done a hands-on activity related to biofilms. Further, 76% (n=45/59) responded that it was easy to obtain supplies for the hands-on activity.

**Figure 4:**
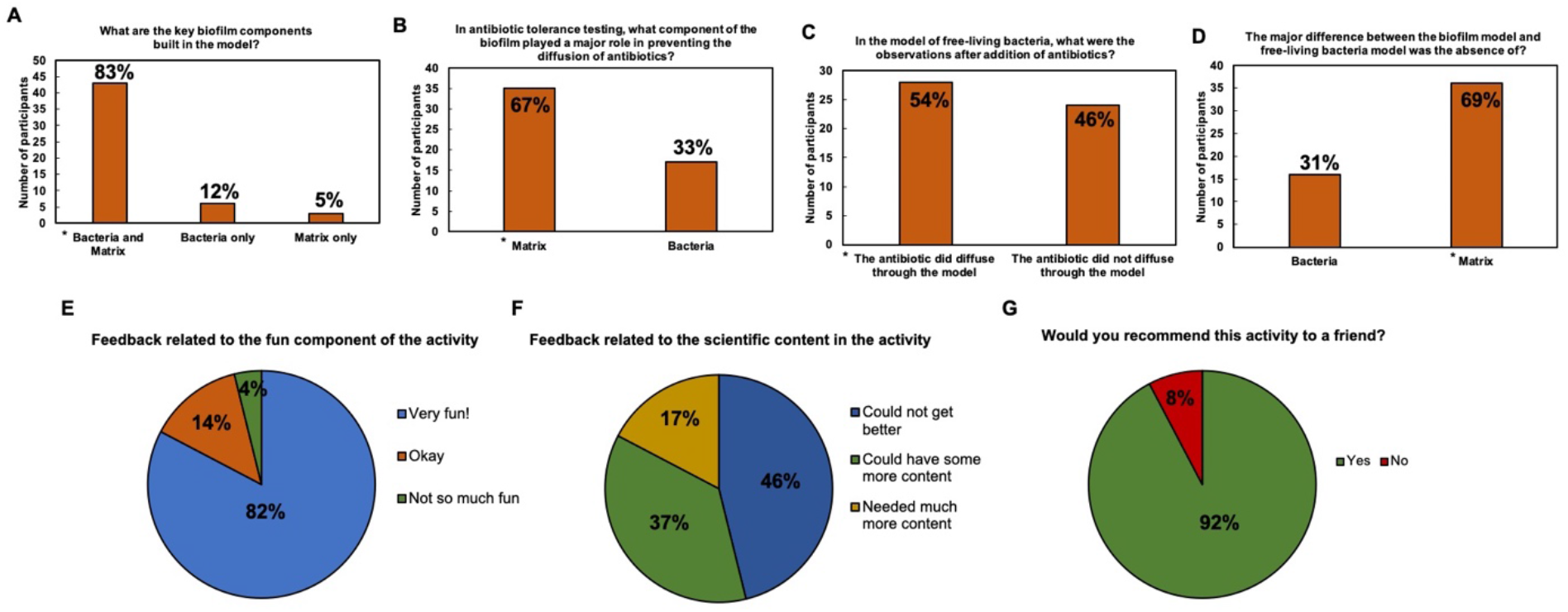
Post-activity feedback from participants with respect learning of key concepts in the hands-on activity and overall experience at the session. **(A)** The majority of the participants, 83% (n=43/52) identified both bacteria and extracellular matrix as the key components used to build the biofilm model. **(B)** In response to the question related to antibiotic tolerance testing, 67% of participants (n=35/52) responded with ‘matrix’ as the major player in the prevention of antibiotic penetration. **(C)** With respect to observations related to the antibiotic response in the model of free-living bacteria, 54% of participants (n=28/52) responded saying that they observed that the antibiotic did penetrate through the model, in contrast to observations with the biofilm model. On the other hand, 46% (n=24/52) responded that they observed antibiotic to not penetrate through the model of free-living bacteria. **(D)** The majority of the participants, 69% (n=36/52) identified the absence of the extracellular matrix as the major difference between the model of free-living bacteria and the biofilm model. **(E)** Based on the fun component of the hands-on activity, 82% of the participants (n=43/52) rated the activity as very fun. **(F)** With respect to scientific content in the activity, 46% (n=24/52) responded that it could not get better, whereas 37% (n=19/52) responded could have some more content. **(G)** The vast majority of the participants, 92% (n=48/52) responded that they would recommend the activity to a friend.

## CONCLUSIONS

‘Biofilms with a dash of color’ is a hands-on activity for school children to build a biofilm model with materials that mimic bacteria and extracellular matrix, following which they use the model to understand the role of the extracellular matrix in antibiotic tolerance in biofilms. The activity is well-suited for delivery in a virtual format and for students to perform at home; the supplies are simple and easy to obtain, and the procedures do not require specialized laboratory equipment. Based on delivery and feedback, the students were able to perform the activity with minimal supervision, comprehend the important concepts related to the observations, and found the session to be informative and fun. Taken together, this means that the activity could be included as part of school science or biology classes, for both in-person and home-based schooling. In doing so, it will serve to understand two fundamental concepts in microbiology: that microbial communities most often exist in self-assembled biofilms, and that the structure of the biofilms contributes to antibiotic tolerance. For students, this could lead to interesting questions and ideas related to the of understanding microbial communities, approaches to overcome antibiotic tolerance in biofilms, and the judicious use of antibiotics.

## SUPPLEMENTAL MATERIALS

**Appendix 1:** Introduction to microbes, including free-living bacterial forms

**Appendix 2:** Introduction to biofilms and antibiotic tolerance in biofilms

**Appendix 3:** Pre-session feedback form

**Appendix 4:** Post-session feedback form

## ACKNOWLEDGEMENTS

Karishma S Kaushik’s academic appointment is supported by the Ramalingaswami Re-entry Fellowship (Department of Biotechnology, Government of India, BT/HRD/16/2006). This virtual science activity was first conducted as part of the Talk to a Scientist (India) science outreach platform (co-founded by KSK and SK), in partnership with India Science Festival (2022). The program Talk to a Scientist (India) is funded by the IndiaBioscience Outreach Grant (IOG) and the American Geophysical Union SciComm grant. We thank the young minds for their participation in the virtual delivery of this activity, and India Science Festival for the opportunity.

