## Appendix 1 for "Biofilms with a dash of color: A hands-on activity for school students to build a biofilm model, and use it to understand antibiotic tolerance in biofilms"

**Introduction to microbes, including free-living bacterial forms**

(Basic content for delivery via slides or chalkboard prior to the hands-on activity)

**1. Introduction to microbes**

Microbes or microorganisms are microscopic living organisms that are invisible to the naked eye, and are therefore viewed under a microscope. They include bacteria, viruses, fungi, archaea and protists. Some examples of bacteria are: *Staphylococcus aureus, Streptococcus, Escherichia coli, Pseudomonas aeruginosa.*

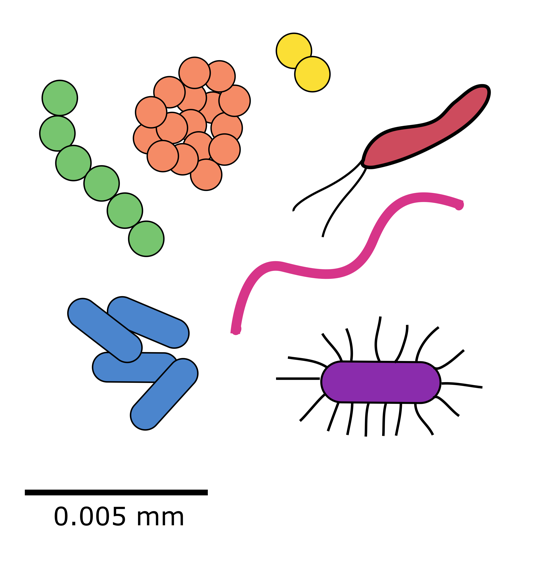

**2. Size and types of bacteria**

Bacteria can vary in size, but typically measure 1-5 micrometers (μm) in size (1 μm = 0.001 mm). They are also observed to have various shapes such as:

**Cocci** (spherical or oval shaped bacteria), that can be arranged in pairs (*Streptococcus pneumoniae*), chains (*Streptococcus* spp.) or clusters (*Staphylococcus aureus*).

**Bacilli** (rod-shaped bacteria), such as *Escherichia coli* and *Salmonella typhi.*

Apart from these, bacteria can also be seen as spiral-shaped, comma-shaped and filamentous forms.

**3. Bacteria in the environment and human body**

Bacteria can be found in the environment and in the human body, where they can have a range of beneficial as well as detrimental effects. In the environment, bacteria play an important role in decomposition, and nitrogen and carbon fixation. In the human body, bacteria such as *Escherichia coli* are normally present in the gastrointestinal tract, and mediate several digestive and immune processes. Bacteria are also causative agents of infections in humans, including a range of eye, ear, skin, lung, gastrointestinal tract, urinary and bloodstream infections.

**4. Bacterial infections are treated with antibiotics**

Bacterial infections are treated with antibiotics; compounds that inhibit important processes in bacteria. This includes the synthesis of important bacterial structures such as the cell wall or processes such as protein synthesis. Given this, antibiotics typically act on actively-growing bacteria, and can either inhibit the growth of bacteria (bacteriostatic) or kill bacteria (bactericidal).

Some examples of antibiotics include: *Penicillin*, *Azithromycin, Tetracycline.*

**Additional reading**

Prescott’s Microbiology. Joanne Willey, Kathleen Sandman, Dorothy Wood. Publisher: McGraw-Hill Higher Education. Print ISBN: 9781260211887, 1260211886, eText ISBN: 9781260409062, 1260409066, Edition: 11^th^, Copyright year: 2020
