## Appendix 2 for "Biofilms with a dash of color: A hands-on activity for school students to build a biofilm model, and use it to understand antibiotic tolerance in biofilms"

**Introduction to biofilms and antibiotic tolerance in biofilms**

(Basic content for delivery via slides or chalkboard prior to the hands-on activity)

**1. What are biofilms?**

Biofilms are aggregates of microbial communities, often bacteria and fungi, that are attached to each other or a surface, and embedded in a self-produced extracellular matrix. The extracellular matrix typically comprises of polysaccharides, proteins, extracellular DNA and water. In the human body, biofilms contribute to a range of infection states, including wound and lung infections. Under these conditions, bacteria in biofilms are tolerant to clearance by the immune cells, as well as treatments with antibiotics.

**2. Proposed reasons for the antibiotic tolerance in biofilms**

Biofilms display tolerance to treatment with most major groups of antibiotics. This is in part due to the structure of biofilms that renders then inaccessible to antibiotics. Additional factors such as dormant bacterial cells as well as enzymes released by bacteria that neutralize antibiotics, also contribute to the antibiotic tolerance of biofilms. Taken, together the antibiotic tolerance of biofilms is a serious health problem, and ongoing research is focused on understanding this better, as well as developing novel approaches to treat biofilms.

**3. The role of the extracellular matrix in the antibiotic tolerance of biofilms**

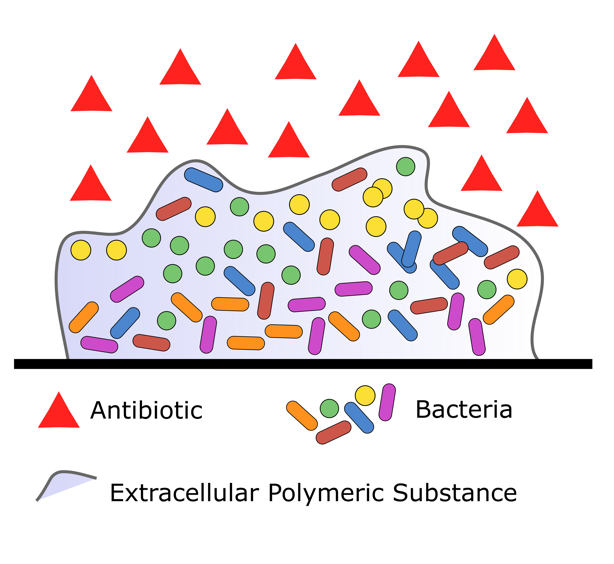
Biofilms consists of aggregates of bacteria, encased in self-produced extracellular matrix. This matrix provides protection to bacteria in the biofilm, transports nutrients and water, as well as provides mechanical support to the biofilm. The extracellular matrix plays an important role in the tolerance on biofilms to antibiotics. This is either via the entrapment or inactivation of antibiotics. Importantly, molecules in the extracellular matrix (polysaccharides, proteins) interact with or entrap antibiotics, and thereby slow or impede the diffusion of antibiotics (known as diffusion-reaction inhibition) through the biofilm. As a result, bacteria in the biofilm are either not exposed to the antibiotic or are exposed to low antibiotic concentrations that are not sufficient to inhibit or kill bacteria.

**Additional Reading**

Watnick P, Kolter R (2000). Biofilm, City of Microbes. *J Bacteriol* 182 (10), 2675-2679.

Flemming H-C, Wingender J, Szewzyk U, Steinberg P, Rice SA, Kjelleberg S (2016). Biofilms: an emergent form of bacterial life. *Nature Reviews Microbiology* 14, 563-575.
