## Appendix 3 for "Biofilms with a dash of color: A hands-on activity for school students to build a biofilm model, and use it to understand antibiotic tolerance in biofilms"

**Pre-activity feedback**

(Feedback questions can be modified depending on location and participant groups)

1. What is your age? (6-10 years / 11-15 years / 16-18 years)

2. What is your education level? (Primary school / Middle School / High school)

3. Where are you joining from? (India / Out of India)

4. Have you heard of biofilms before? (Yes / No)

5. Have you done a hands-on activity on biofilms before? (Yes / No)

6. Was it easy to obtain supplies for today’s hands on activity? (Yes / No)
