## Appendix 4 for "Biofilms with a dash of color: A hands-on activity for school students to build a biofilm model, and use it to understand antibiotic tolerance in biofilms"

**Post-activity feedback**

1. What are the major biofilm components built in the model?

(Bacteria only / Bacteria and matrix / Matrix only)

2. In antibiotic tolerance testing, what component of the biofilm played a key role in preventing the diffusion of antibiotics?

(Bacteria / Matrix)

3. In the model of free-living bacteria, what were the observations after addition of antibiotics?

(The antibiotic did diffuse through the model / The antibiotic did not diffuse through the model)

4. The major difference between the biofilm model and free-living model was the absence of?

(Bacteria / Matrix)

5. How would you rate the activity based on fun?

(Very fun / Okay / Not so fun)

6. How would you rate the activity based on content?

(Could not get better / Could have some more content / Needed much more content)

7. Would you recommend this hands-on activity to a friend? (Yes/No)
